# RiboTag RNA Sequencing Identifies Local Translation of HSP70 In Astrocyte Endfeet After Cerebral Ischemia

**DOI:** 10.1101/2024.10.08.617236

**Authors:** Bosung Shim, Prajwal Ciryam, Cigdem Tosun, Riccardo Serra, Natalia Tsymbalyuk, Kaspar Keledjian, Volodymyr Gerzanich, J. Marc Simard

## Abstract

Brain ischemia causes disruption in cerebral blood flow and blood-brain barrier (BBB) integrity which are normally maintained by the astrocyte endfeet. Emerging evidence points to dysregulation of the astrocyte translatome during ischemia, but its effects on the endfoot translatome are unknown. In this study, we aimed to investigate the early effects of ischemia on the astrocyte endfoot translatome in a rodent model of cerebral ischemia-reperfusion. To do so, we immunoprecipitated astrocyte-specific tagged ribosomes (RiboTag IP) from mechanically isolated brain microvessels. In mice subjected to middle cerebral artery occlusion and reperfusion and contralateral controls, we sequenced ribosome-bound RNAs from perivascular astrocyte endfeet and identified 205 genes that were differentially expressed in the translatome after ischemia. Pathways associated with the differential expressions included proteostasis, inflammation, cell cycle, and metabolism. Transcription factors whose targets were enriched amongst upregulated translating genes included HSF1, the master regulator of the heat shock response. The most highly upregulated genes in the translatome were HSF1-dependent *Hspa1a* and *Hspa1b*, which encode the inducible HSP70. We found that HSP70 is upregulated in astrocyte endfeet after ischemia, coinciding with an increase in ubiquitination across the proteome. These findings suggest a robust proteostasis response to proteotoxic stress in the endfoot translatome after ischemia. Modulating proteostasis in endfeet may be a strategy to preserve endfeet function and BBB integrity after ischemic stroke.

## Introduction

Ischemic stroke remains a leading cause of death in the US and worldwide [1–3]. The vast majority of strokes are caused by cerebral ischemia, which results in cerebral edema [4], impaired cerebral blood flow (CBF) [5], neuroinflammation [6], blood-brain barrier (BBB) disruption [7], and widespread cell death [8]. Astrocytes play a central role in maintaining an intact central nervous system (CNS) and responding to brain injury. In particular, astrocytes are responsible for maintaining BBB integrity [9], regulating cerebral blood flow [10], and locally transporting ions and metabolites [11,12]. Many of these functions occur at the perivascular endfeet, which directly contact and ensheathe cerebral blood vessels as part of the neurogliovascular unit [13]. As such, astrocytes and their perivascular endfeet are important potential therapeutic targets for improving outcomes in ischemic stroke.

Perivascular endfeet are located long distances from the cell body and possess the necessary machinery, including the endoplasmic reticulum and Golgi apparatus [14], to sustain a specialized proteome [15,16]. Recently, protein translation of a large repertoire of endfoot-localized mRNA has been reported. Similarly, local translatomes have been identified in perisynaptic processes [17,18]. These seminal findings were made following the conception and optimization of the RiboTag approach to isolate ribosome-bound RNA from specific brain cell types from RiboTag transgenic mice [19,20].

There is growing evidence of major shifts in the astrocyte translatome acutely after ischemia, including the translation of stress response genes and transcription factor [21,22]. It is not known whether a distinct translational response occurs specifically in the astrocyte endfeet in response to ischemia. In the present study, we aimed to characterize the translatome of perivascular endfeet at an early reperfusion timepoint in mice following severe ischemia.

## Materials and Methods

### Mice and Surgical Procedure

Astrocyte-specific RiboTag mice (FVB-Tg(Aldh1l1-EGFP/Rpl10a)JD133Htz/J, #030248, Jackson Laboratory) were bred and maintained in-house (University of Maryland School of Medicine). Transgene expression was confirmed by genotyping using collected tail samples (Transnetyx). Surgery to induce middle cerebral artery occlusion (MCAo) in male (10-12 week old) mice was performed as previously described [23], that is common carotid artery ligation and intraluminal occlusion using a 6-0 silicon filament occluder (Doccol Corp). Following placement of the occluder, MCAo animals recorded > 70% reduction in relative cerebral blood flow (rCBF) as measured using laser Doppler flowmetry (DRT2, Moor Instruments). Following 2- hour occlusion, the filament occluder was removed and the mice were allowed to recover.

### Brain Microvessel and Astrocyte Endfeet Isolation

Following 6-hours of reperfusion, mice were terminally anesthetized by pentobarbital overdose then transcardially perfused with ice-cold normal saline (0.9%). MCA territories were dissected then homogenized by hemisphere in a 7-mL glass Dounce homogenizer (12-15 strokes) in 1× HBSS containing Ca2+/Mg2+ (Gibco) and supplemented with HEPES. Homogenates were centrifuged at 2000×G for 10min at 4°C. Pellets were resuspended in 18% dextran (from Leuconostoc spp., M_r_∼70000, MilliporeSigma) then centrifuged at 10000×G for 15min at 4 °C. Myelin-cleared pellets were resuspended in 1% bovine serum albumin then filtered through a 20-um nylon mesh. Microvessels caught in the mesh were retrieved and centrifuged at 2000×G for 10min. For immunostaining, microvessels were cytospun onto glass microscope slides and fixed in 4% paraformaldehyde for 10min at 4°C prior to storage in -20°C. For RiboTag immunoprecipitation, solutions were supplemented with 100ug/ml cycloheximide and 200U/ml RNAsin. Astrocyte endfeet isolation were performed as previously described [15]. Briefly, isolated microvessels were dissociated into single cells following incubation in 100µg/ml Liberase DL (MilliporeSigma) at 37°C for 30min. Larger cells and debris were removed by centrifugation at 300×G at 4°C for 30min. The supernatant was centrifuged at 25000×G at 4°C for 30min, and the pellet was resuspended in AstroMACS separation buffer (Miltenyi). Following labelling with magnetic bead-conjugated anti-ACSA2 (Miltenyi), ACSA2+ astrocyte endfeet were positively selected and immediately processed for protein analysis.

### RiboTag Immunoprecipitation (IP) and RNA-seq

For RiboTag IP, all steps were performed in 4°C and RNase-free (RNaseZAP, ThermoFisher) conditions. Microvessels were lysed in freshly prepared lysis buffer (150mM KCl, 20mM HEPES [pH 7.4), 5mM MgCl2, 1mM DTT, 1% NP-40, 100ug/ml cycloheximide, 30mM DHPC [1,2-dicaproyl-sn-glycero-3-phosphocholine, 07:0]) supplemented with 200U/ml RNAsin (ProMega) and cOmplete protease inhibitor (MilliporeSigma). Following homogenization using a handheld pestle motor mixer, lysates were centrifuged at 18000×G in 4°C for 10min. IP was performed by incubating the soluble fraction with magnetic bead-conjugated anti-GFP antibody (#67090, Cell Signaling Technology) overnight (∼16 hours) with end-over-end rotation. Following magnetic separation from the supernatant (DynaMag-2, ThermoFisher), beads were washed three times in a high-salt (300mM KCl) lysis buffer supplemented with 200U/mL RNAsin. Unpurified RNA was collected from the magnetic beads using the TriZOL reagent according to the manufacturer’s instructions. Total mRNA (in ultrapure water) following RiboTag IP was quantified using the Bioanalyzer 2100 (Agilent). Strand-specific RNA sequencing was performed by Maryland Genomics in NovaSeq S4 flow cells (NovaSeq6000, Illumina) at 100bp paired end reads with 67 ± 18 million (mean ± SD) reads per sample.

### Sequence Alignment

Raw R1 and R2 fastq files were trimmed to remove adapter sequences and low quality reads using the Trimmomatic software v0.33 under the following parameters: simple clip threshold (7), seed mismatches (2), palindrome threshold (40), minimum sequence length (3), and trailing quality (20). Trimmed sequences were followed by quality control evaluation using FastQC. Trimmed sequences were aligned using STAR v2.7.5a [24] to the *Mus musculus* mm39 primary assembly. These were quantified at the gene level using the STAR geneCounts tool to obtain reverse/forward strand-specific counts. Analysis of unmapped, multi-mapped, and ambiguously mapped reads was performed using blastn [25] with default settings.

### Differential Gene Expression Analysis

Analysis was performed using DESeq2 v1.44.0 [26] in R (v4.4.1). There were 10,862 genes that had ≥ 10 counts per sample in ≥ 3 samples. We assessed the pairwise correlation of gene counts in these samples after rlog transformation. We generated a model for differential expression analysis using two variables: 1) individual animal (ID 1, 2, or 3) and 2) disease state (Control or Stroke). We performed fold change shrinkage using the adaptive shrinkage method ashr. Significance testing was performed against the null hypothesis that |log2(Fold Change)| < 0.322, known as the *s*-value, corresponding to gene expression in the ischemia/reperfusion condition of <4/5 or >5/4 that of controls [27].

### Principal Component Analysis (PCA)

PCA of the gene counts was performed using the R package PCAtools v2.16.0. The correlation between the first 6 principal components and the sample variables was performed using the eigencorplot function. We generated the biplot as described in PCAtools, plotting each sample using the centered input data multiplied by the variable loadings (i.e., “rotated data”) for principal components 1 and 2. Rescaled loadings are also plotted. The scaled loading for a given gene (scaledx) equal to the loading value for that gene (loadingx) by 1.5 times the minimum value of the ratio of the range of the rotated data to the range of the loadings data for principal component:

### Gene Set Enrichment Analysis (GSEA)

GSEA was performed using the R package fgsea v1.30.0 [28–30]. Three sets of pathway databases were obtained from the mouse Molecular Signatures Database [31]: hallmark gene set (MH, v2024.1), canonical pathway curated gene set (M2.CP, v2024.1), and Gene Ontology (GO) biological processes gene set (M5.GO.BP, v2024.1). To compare pathways to our gene set, a mapping table was generated of Ensembl gene IDs to gene symbols by automated conversion supplemented with manual curation. A pathway was included in enrichment analysis if our gene expression data set included at least 10 genes from that pathway and 50% of total genes in the pathway. This resulted in 44/50 MH, 965/1730 M2.CP, and 3719/7713 pathways included in downstream analysis. Enrichment was performed using the fgsea function fgseaMultilevel, which uses the adaptive multilevel splitting Monte Carlo approach, with default settings. Ranking of genes for enrichment analysis was done in one of two ways: based on 1) fold change or 2) the Wald statistic, and a pathway significance threshold was set at Benjamini-Hochberg adjusted p < 0.00833 (≈0.05/6 to account for multiple testing across 3 pathway gene sets with 2 ranking approaches). Consensus enriched pathways were identified that were significant by both the fold change and Wald statistic metrics. Pathways were then manually annotated into categories as shown in Figure 2.

### Gene Structural Analysis

From the GENCODE protein-coding transcript sequences file, sequences and annotations for 5’-UTR and 3’-UTR sequences were extracted.

### Transcription Factor Target Enrichment Analysis

Transcription factor and target mapping were obtained from the TFLink database (TFLink_Mus_musculus_interactions_All_GMT_ncbiGeneID_v1.0.gmt) [32]. To compare transcription factor target sets to our gene set, a mapping table was generated of Ensembl gene IDs to NCBI gene IDs by automated conversion supplemented with manual curation. Enrichment analysis was performed using Fisher’s exact test with multiple hypothesis correction performed using the Benjamini-Hochberg method.

### Immunohistochemistry (IHC), Fluorescence Imaging, and Stimulated Emission Depletion (STED) Imaging

After terminal anesthesia using excess pentobarbital, mice were transcardially perfused with ice-cold normal saline (0.9%) then fixed with 4% paraformaldehyde. Whole brains were dissected, post-fixed overnight and cryopreserved in 30% sucrose (in 1× PBS) for at least 48 hours. 12-um thick coronal brain sections were prepared on glass microscope slides by cryosectioning. Tissue sections or isolated microvessels on glass microscope slides were immunostained as previously described [15]. Primary and fluorophore-conjugated secondary antibodies are listed in Table S1. Slides were coverslipped using either ProLong Gold Antifade mount (Invitrogen) for fluorescence imaging or the liquid antifade mountant (Abberior) for STED imaging. Omission of primary antibody was included to verify specificity of the immunolabelling. Fluorescent images were acquired using an upright NiE fluorescence microscope system (Nikon) accompanied with the 20× (Nikon Plan Apo λ 20× / 0.75 NA, WD 1.0mm) or 100× (Nikon Plan Apo λD 100× / 1.45oil NA) objectives. Excitation and emission wavelengths were as follows (in nm): 488/525, 561/620, and 640/710. Images were processed and analyzed using the NIS Elements v5.30 (Nikon). STED imaging was performed using the STED Facility Line microscopy system (Abberior). Raw STED images were acquired using the 60× (Olympus UPlanSApo 60× / 1.30sil NA) objective. Signal for STED imaging was excited at 640nm with pulsed depletion at 775nm. Images were acquired using the Imspector Image Acquisition software and further processed on ImageJ v1.54f (Fiji). Both fluorescent and STED images were acquired at equal laser strength and exposure times between non-ischemic control and ischemic conditions.

### Quantitative reverse-transcription PCR (qPCR) of Translating Endfeet RNA

Following RiboTag IP and RNA extraction, RNA amount was quantified using the Qubit 4 Fluorometer (ThermoFisher). First-strand cDNA was generated from 40ng RNA using the SuperScript IV reverse transcription kit (ThermoFisher) according to the manufacturer’s recommendations. qPCR reactions were prepared using the PowerUp SYBR Green master mix with (per reaction) 1 µl cDNA and 200 nM each of the forward and reverse primers. qPCR was performed using the ABI7300 (Applied Biosystems) with the following thermal cycling conditions: 50°C for 2min, 95oC for 2min, then 40 cycles of 95oC for 15sec and 60°C for 30sec. Melting curve analysis was included in all qPCR reactions to confirm that a single product was amplified per reaction. Primers (sequences listed in Table S2) were designed to span exon-exon junctions. Raw CT values were normalized to that of Gapdh for each biological sample, then quantified relative to the mean of the fold changes of microvessels or non-stroke control mRNA.

### Immunoblotting

Protein was extracted from brain homogenates and isolates using 1× RIPA buffer supplemented with cOmplete protease inhibitor (MilliporeSigma). Following homogenization using a handheld pestle motor mixer, samples were centrifuged at 18000×G in 4°C for 10min. Supernatant containing soluble proteins were separated from the pelleted insoluble fraction. Protein amount was quantified using the Qubit 4 fluorometer and Qubit protein assay kit (ThermoFisher). Lysates were denatured at 70°C for 5min in NuPAGE LDS sample buffer and sample reducing agent (ThemoFisher). Conventional protein gel electrophoresis, blot transfer, and immunoblotting (in 5% nonfat milk in 1× TBST) were performed. Primary and secondary antibodies used for immunoblotting are listed in Table S1. Blots were visualized using the SuperSignal West Atto Ultimate Sensitivity substrate (ThermoFisher). Densitometry analysis was performed using ImageJ v1.54f (Fiji).

### Statistical Analysis and Figure Preparation

Unless specified otherwise, data are presented as mean ± SEM and the specific analysis are included in the figure legends. Figures were prepared using NIS Elements (Nikon), ImageJ (Fiji), and R including ggplot2 (R Foundation). Schematics were created using BioRender.com.

## Results

### Validation of Endfeet RiboTag Isolations after MCAO

To characterize the astrocyte endfoot-specific translatome from the post-ischemic brain, we immunoprecipitated RiboTag and ribosome-bound RNA from mechanically isolated microvessels [33], as previously reported (Figure 1A) [14]. The RiboTag IP relies on the transgenic expression of a tagged Rpl10a ribosomal protein driven by an Aldh1l1 promoter, which largely restricts expression to astrocytes [14,20]. Astrocyte specificity has been shown to persist following brain ischemia [22].

**Figure 1.**
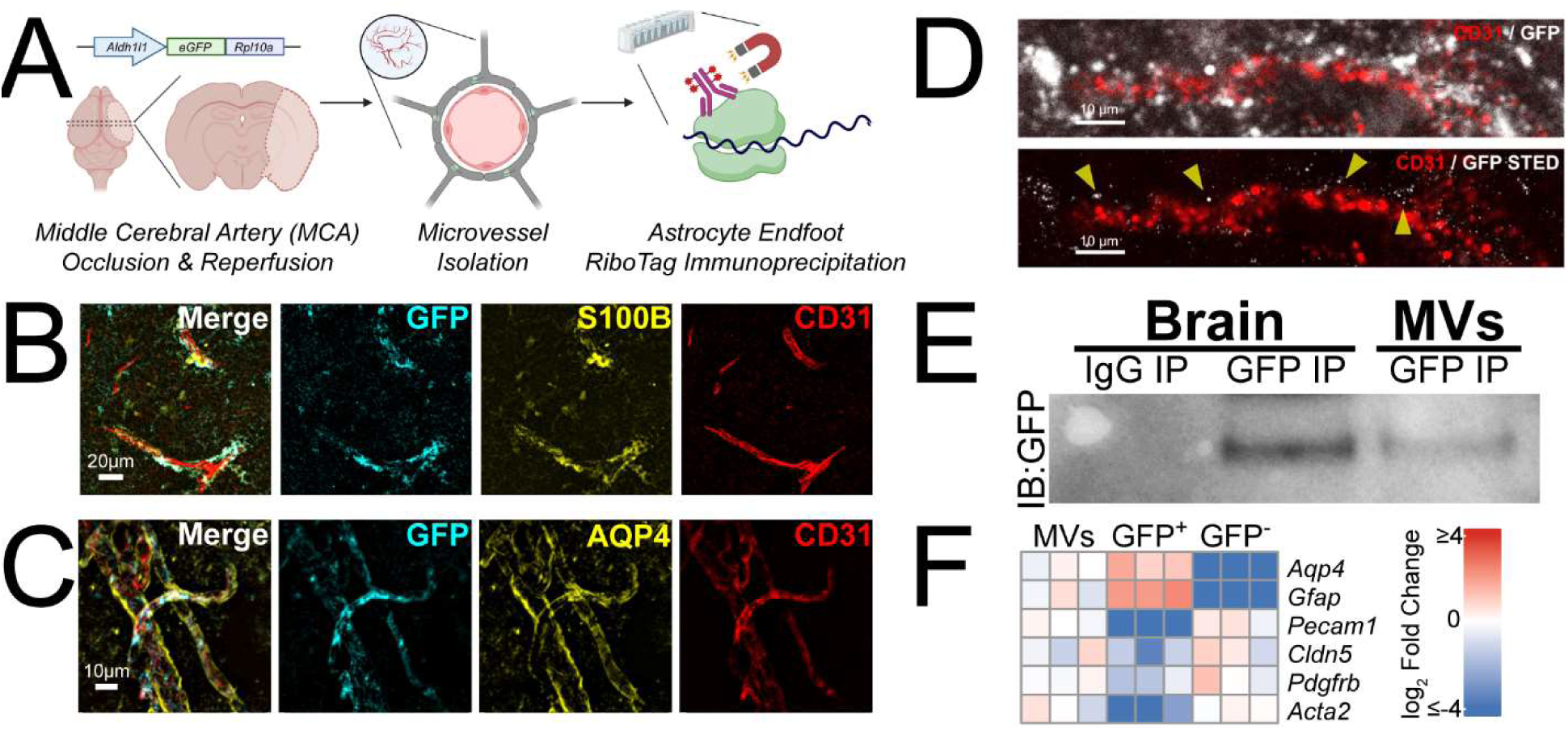
Detection of RiboTag and ribosome-bound mRNA from post-ischemic astrocyte endfeet. (A) Schematic illustrating the induction of stroke via transient MCAo/R, followed by isolation of microvessels from the post-ischemic brain tissue and RiboTag IP. (B) Representative high-powered image of the post-ischemic cortex with immunolabelling for GFP (cyan), S100B (yellow), and CD31 (red) (scale bar = 20µm). (C) Representative image of microvessels mechanically isolated from the post-ischemic tissue with immunolabelling for GFP (cyan), AQP4 (yellow), and CD31 (red) (scale bar = 10µm). (D) High-powered images without (*above*) or with (*below*) STED of isolated post-ischemic microvessels immunolabelled for GFP (white) and CD31 (red) (scale bar = 10µm). (E) Representative immunoblot following RiboTag IP from post-ischemic brain lysate or microvessels. (F) qPCR of post-ischemic microvessels before RiboTag IP (MVs), after RiboTag IP (GFP+), or non-IP’d flowthrough (GFP-). Data expressed as mean ± SEM (n = 3 biological replicates).

We first sought to validate the approach for use in mice following 2 hours ischemia via MCAo and 6 hours reperfusion, given that ischemia induces morphological changes of perivascular endfeet such as swelling [34] and degeneration [35]. We performed IHC in brain sections of GFP in the transgenic mice following MCAo and found that GFP was detected in S100B+ astrocytes, including in perivascular domains with minimal localization over CD31+ vessels (Figure 1B). The GFP signals were detected in isolated microvessels from the post-ischemic brain including co-labelling over AQP4+ astrocyte endfeet, suggesting that RiboTag expression survives the mechanical isolation of microvessels (Figure 1C). STED imaging to visualize nanometer-range GFP puncta revealed that GFP remains primarily perivascular (Figure 1D). Immunoblotting for GFP following RiboTag IP from post-ischemic microvessel lysates confirmed detectable RiboTag expression (Figure 1E). qPCR of RiboTag immunoprecipitated mRNA from isolated microvessels after stroke revealed enrichment of astrocyte markers Aqp4 and Gfap [36,37] as well as depletion of vascular markers including Pecam1, Cldn5 (endothelial cells), Pdgfrb (pericytes), and Acta2 (vascular smooth muscle cells) (Figure 1F). These results collectively verified that perivascular astrocytic RiboTag and ribosome-bound mRNA were detectable in post-ischemic isolated microvessels.

### Quality assessment of RiboTag RNA sequencing

To identify changes in the translatome induced by stroke, we extracted RNA from endfeet ribosomes and performed RNA sequencing. As it has been reported that poly(A) selection can introduce bias into RNA sequencing data sets [38], we sought to determine whether adequate sequencing depth could be achieved without selection. As it has been reported that poly(A) selection can introduce bias into RNA sequencing data sets [38], we sought to determine whether adequate sequencing depth could be achieved without selection. We sequenced 67 ± 18 million reads per sample, of which 63 ± 1% were mapped uniquely. Of those reads not uniquely mapped, we aligned 1 million per sample against the genomic sequence of the *Mus musculus* 45S pre-ribosomal RNA (NR_046233.2) and determined that 82 ± 0.02% were ribosomal in origin (Figure 2A). This suggests that the unique mapping rate is a result of the absence of complete rRNA depletion. As the proportion of rRNA is expected to be much higher in whole transcriptome samples, our results suggest that the RiboTag approach substantially enriches for mRNA, which may serve as an internal control for the quality of ribosome isolation. As another quality control measure, we found read counts to be > 90% correlated with each other across all samples (Figure 2B).

**Figure 2.**
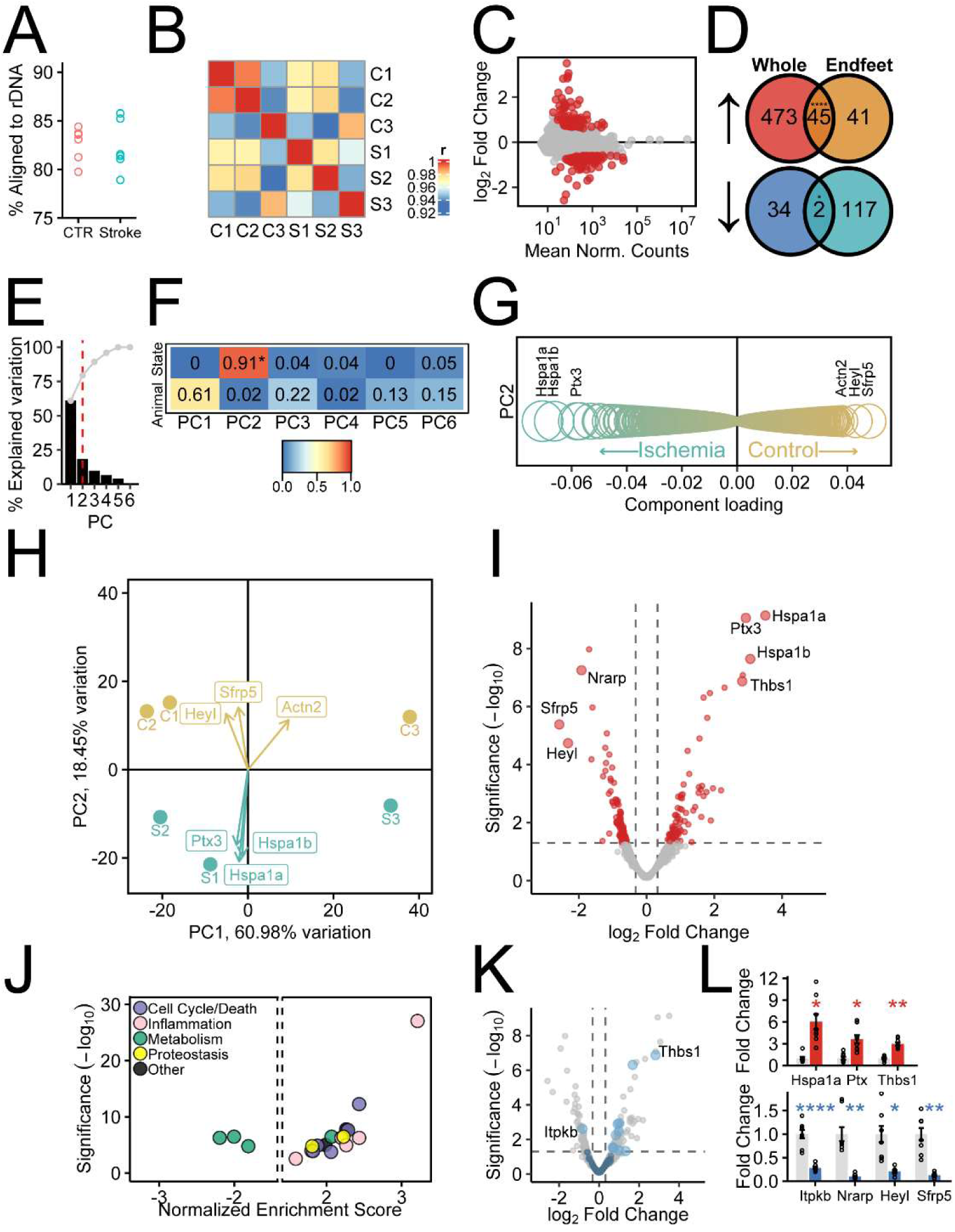
RNA sequencing analysis of ribosome-bound RNA from post-ischemic astrocyte endfeet. (A) Percentage of reads not uniquely mapped to the mouse genome that were subsequently mapped to mouse ribosomal DNA. (B) Correlation matrix of the entire sample set (3 contralaterals C1-C3, 3 stroke S1-S3) used in this study. Pearson *r* was used for the heatmap color scheme. (C) MA scatterplot showing the relationship of fold change to mean normalized counts after log fold change shrinkage. Red points are differentially expressed. (D) Venn diagrams of the 205 differentially expressed endfoot genes (86 upregulated, 119 downregulated) after ischemia plotted against DEGs identified in an existing astrocyte translatome during the hyperacute phases of ischemia [22]. Significance of enrichment tested using the Fisher exact test, with Holm-Bonferroni adjustment. (E) Scree plot of principal components (PCs) with threshold at PC2 indicated by the red dashed line. (F) Correlation matrix plotted for the tested variables (Disease State or Animal) against the six PCs, with Holm-Bonferroni adjusted Pearson *r*^2^ displayed. (G) Component gene loadings for PC2, with relative loading values depicted by the size of the circles. (H) Principal component biplot showing individual endfoot samples C1-C3 and S1-S3 overlayed with the most negative and positive gene loadings contributing to variation explained by PC2. (I) Volcano plots depicting changes in post-ischemic endfoot gene expression. Vertical grey lines show threshold of log_2_Fold Change significance (±0.322). Grey line shows threshold of significance (*s*-value < 0.05). (J) Significance (Benjamini-Hochberg adjusted *p*-values) and normalized enrichment score from GSEA analysis for Hallmark pathways found to be enriched in upregulated or downregulated genes based on both fold change and Wald statistic metrics. (K) Volcano plot of as in (H), highlighting calcium-regulated genes. (L) qPCR validation of the most upregulated (*Hspa1a*, *Ptx3*, *Thbs1*) and downregulated (*Sfrp5*, *Heyl*, *Nrarp*, *Itpkb*) genes. Relative fold changes per gene normalized to *Gapdh* are expressed as mean ± SEM (\**p* < 0.05, \*\**p* < 0.01, \*\*\*\**p* < 0.0001).

### Features of the Post-Ischemic Endfoot Translatome

Principal component analysis of the transcriptome data set revealed two major components contributing to the variation in the data (Figure 2E-G, Table S3). Contralateral (CTR) and ischemia/reperfusion (Stroke) samples were clearly distinguished by PC2 (Figure 2F-H). From our dataset, we found that 86 genes were upregulated, and 119 genes were downregulated in post-ischemic endfeet compared to contralateral endfeet. Of the six genes that contribute most to the variation in PC2, five are significantly differentially expressed: *Hspa1a* (log2Fold Change: 3.51, *s*-value: 1.37×10^-08^), *Hspa1b* (log2Fold Change: 3.06, *s*-value: 3.21×10^-07^), *Ptx3* (log2Fold Change: 2.93, *s*-value: 2.72×10^-08^), *Sfrp5* (log2Fold Change: -2.58, *s*-value: 4.51×10^-05^), and *Heyl* (log2Fold Change: -2.32, *s*-value: 1.65×10^-04^) (Figure 2F-H, Table S4). We also detected the upregulation of three mitochondrial genes (*mt-Cytb, mt-Nd2,* and *mt-Nd1*), which cytosolic ribosomes would not typically translate. This may reflect the release of mitochondrial transcripts from fragmented mitochondria in the setting of ischemia [39].

We next asked whether the translatome response to stroke in the astrocyte endfeet resembles that previously reported in whole astrocytes in a model similar, but not identical, to our own [22]. To maintain consistency, we re-aligned and analyzed raw data files from RiboTag sequencing reads of astrocytes at four hours after ischemia. We found a highly significant overlap between differentially expressed genes (DEGs) in endfeet and whole astrocytes, especially among upregulated genes (Figure 2D). This included a prominent activation of heat shock response genes like *Hspa1a* and *Hspa1b*, as well as hypoxia-response genes like *Hmox1* and acute phase response genes like Ptx3. Nevertheless, 45% of genes detected as upregulated in endfeet were not detected as upregulated in whole astrocytes. Only two downregulated genes overlapped between the two data sets, the *Hes5* transcription factor gene and the poorly characterized gene *RP23-391P21.4*. As we detect transcription factors and somatic marker genes in our data set, we cannot completely exclude the presence of RNA from cell bodies.

GSEA revealed a range of enriched pathways associated with cell cycle/death, inflammation, metabolism, and proteostasis enriched (Figure 2J, Table S5). Amongst downregulated genes, all enriched pathways were related to metabolism, including oxidative phosphorylation and two pathways related to lipid metabolism (bile acid metabolism and adipogenesis). Notably, the most enriched pathway is the pathway for TNFα signaling via NF-κB. Pathways for the unfolded protein response and the mTOR pathway were also significantly enriched among upregulated genes.

As we recently showed marked calcium influx in perivascular astrocyte endfeet induced by ischemia [23], we wondered whether calcium-responsive genes were differentially expressed in stroke (Figure 2K, Table S6). The Gene Ontology Biological Process term ‘Cellular Response to Calcium Ion’ term was found to be enriched in upregulated genes by (Benjamini-Hochberg adjusted *p*=3.3×10^-2^, NES=1.7) and is the only enriched ion-response term (Table S5). The calcium-regulated transcriptome was recently defined in glomerular podocytes based on calcium ionophore challenge [40]. Like astrocytes, glomerular podocytes participate in regulating vascular permeability [41]. We found no overlap in downregulated genes, but a significant overlap in upregulated genes (odds ratio=9.7, Holm-Bonferroni adjusted Fisher Exact Test *p*-value= *p*=2.7×10^-4^).

Given the prominent role that the heat shock response plays in post-ischemic astrocyte endfoot translatome, we sought to further validate these changes using qPCR (Figure 2L). In addition to the upregulation of *Hspa1a* and *Hspa1b*, we find that *Sfrp5*, the most downregulated gene in our translatome data set, is also decreased in expression based on qPCR. Similarly, the most upregulated and downregulated calcium-responsive genes in *Thbs1* and *Itpkb*, respectively, are also significantly changed in expression in our data set.

### Regulatory features of differentially expressed genes

We next investigated potential mechanisms of regulation of the endfoot translatome. Given recent evidence that RNA structural features are associated with expression in the endfoot translatome, we tested whether sequence length and GC content differed in control and stroke tissue (Figure 3A,B). We found that upregulated genes tend to have longer 5’-UTRs and protein-coding sequences, while downregulated genes tend to have shorter 3’-UTRs and protein-coding sequences. On average, both upregulated and downregulated genes have higher GC content in the protein-coding region than do genes in the translatome as a whole, but only downregulated genes tend to have a higher GC content in the 3’-UTR. Reasoning that these features may affect translation in part through select by specific RNA binding proteins, we tested the association between differential expression in stroke and dependence on the Quaking RNA binding protein (QKI). It was recently shown that deletion of QKI alters the translatome of maturing astrocytes [42]. We found that 9.2% of downregulated genes (odds ratio=10.7, Holm-Bonferroni adjusted Fisher Exact Test *p*-value= *p*= 3.2×10^-8^), but no upregulated genes, were QKI-dependent (Figure 3C). Conversely, an unbiased search of transcription factors using the TFLink database [32] revealed only transcription factors enriched in upregulated genes (Figure 3D, Table S7-S8). These included HSF1, the master regulator of the heat shock response, as well as a number of transcription factors associated with inflammation (e.g., NFκB1) and cell cycle/death (e.g., FOSL1, RB1). Interestingly, DAND5 has been reported to be a negative regulator of angiogenesis [43]. These transcription factors form a network of shared regulation, with many genes under the influence of multiple transcription factors (Figure 3E). While *Hspa1a* and *Hspa1b*, canonical genes in the heat shock response, are transcriptionally activated by HSF1, they differ in terms of additional genetic interactions with other transcription factors. FOSL1, for example, regulates *Hspa1a* but not *Hspa1b*, while DAND5 and SP3 regulate *Hspa1b* but not *Hspa1a*.

**Figure 3.**
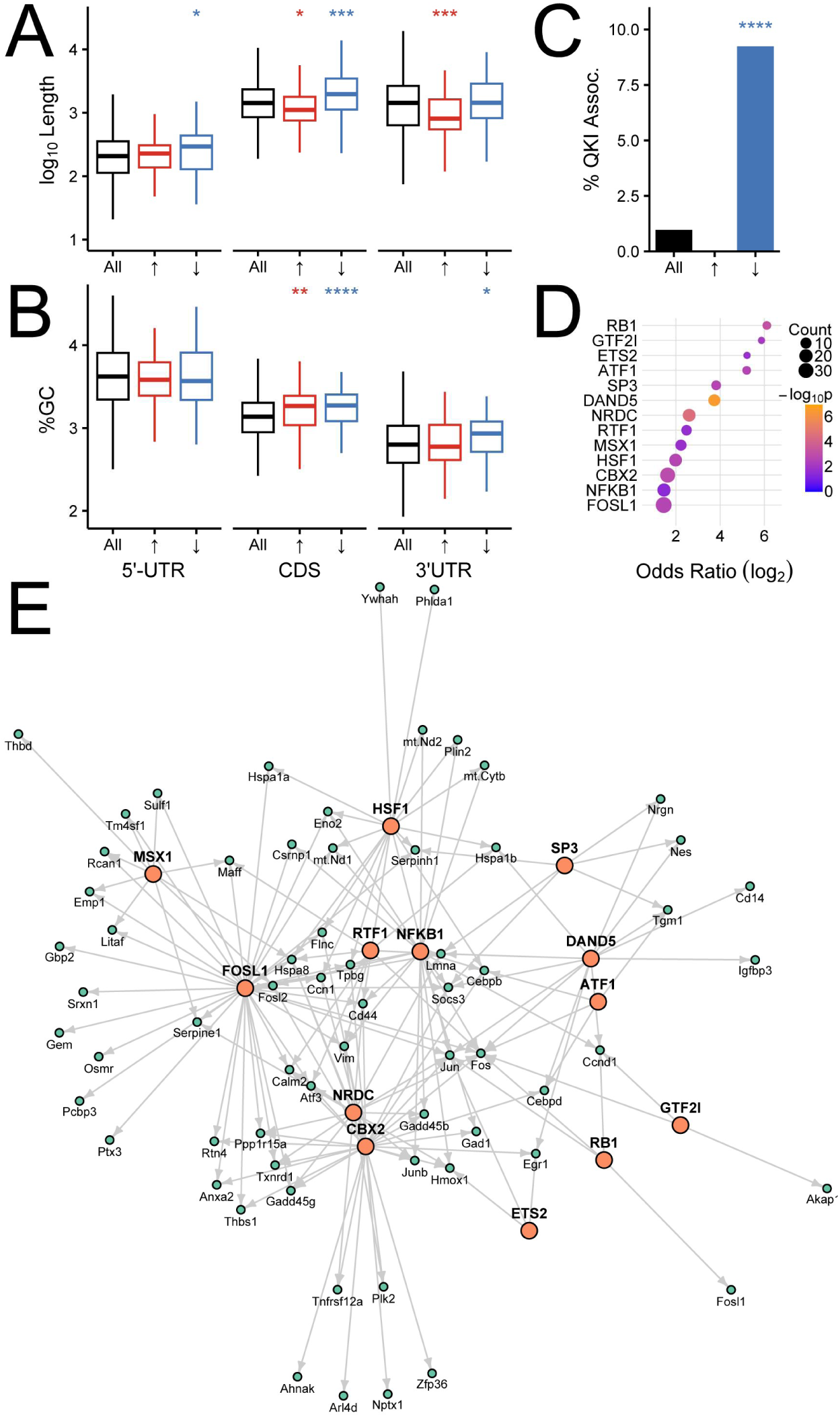
Mechanisms of regulation of the astrocyte endfoot translatome. (A) Length and (B) GC content of the translatome (black), upregulated genes (red), and downregulated genes. Statistical significance was determined using the Wilcoxon-Mann Whitney test with Holm-Bonferroni adjustment. (C) Percent of the translatome (black), upregulated (red, 0%, not visualized), and downregulated (blue) genes that are known to be dependent on Quaking RNA binding protein [42]. (D) Transcription factors enriched in upregulated genes relative to whole translatome. (E) Network diagram of transcription factor and gene target interactions generated with R package igraph using the Fruchterman-Reingold layout algorithm. (\**p* < 0.05, \*\**p* < 0.01, *** *p* < 0.001, \*\*\*\**p* < 0.0001).

### Upregulation of HSP70 and protein ubiquitination in post-ischemic endfeet

Given that *Hspa1a* and *Hspa1b* were the most upregulated genes in the post-ischemic endfoot translatome, we asked whether their protein product, inducible HSP70, was more abundant in endfeet after stroke. We isolated astrocyte endfeet from post-ischemic or control microvessels following proteolytic dissociation to single cells and positively selecting for ACSA2+ endfeet by magnetic separation (Figure 4A). Immunoblotting for HSP70 revealed its upregulation in astrocyte endfeet after ischemia (Figure 4A,B). The constitutively expressed heat shock cognate HSC70, whose corresponding gene was more modestly upregulated in the stroke endfoot translatome, shows a small but significant upregulation in post-ischemic endfeet (Figure 4A,B). Immunolabelling of HSP70 corroborated the upregulation of HSP70 found in immunoblotting, with increased HSP70 immunoreactivity in AQP4-positive endfeet after stroke (Figure 4C). We hypothesized that the robust upregulation of HSP70 and other proteostasis components may have been in response to proteotoxic stress induced by ischemia. Consistent with this, we note a global increase in ubiquitination in stroke (Figure 4D). These results suggest active translation of *Hspa1a* and *Hspa1b* in the astrocyte endfeet after stroke are likely in response to global disruptions in endfoot proteostasis.

**Figure 4.**
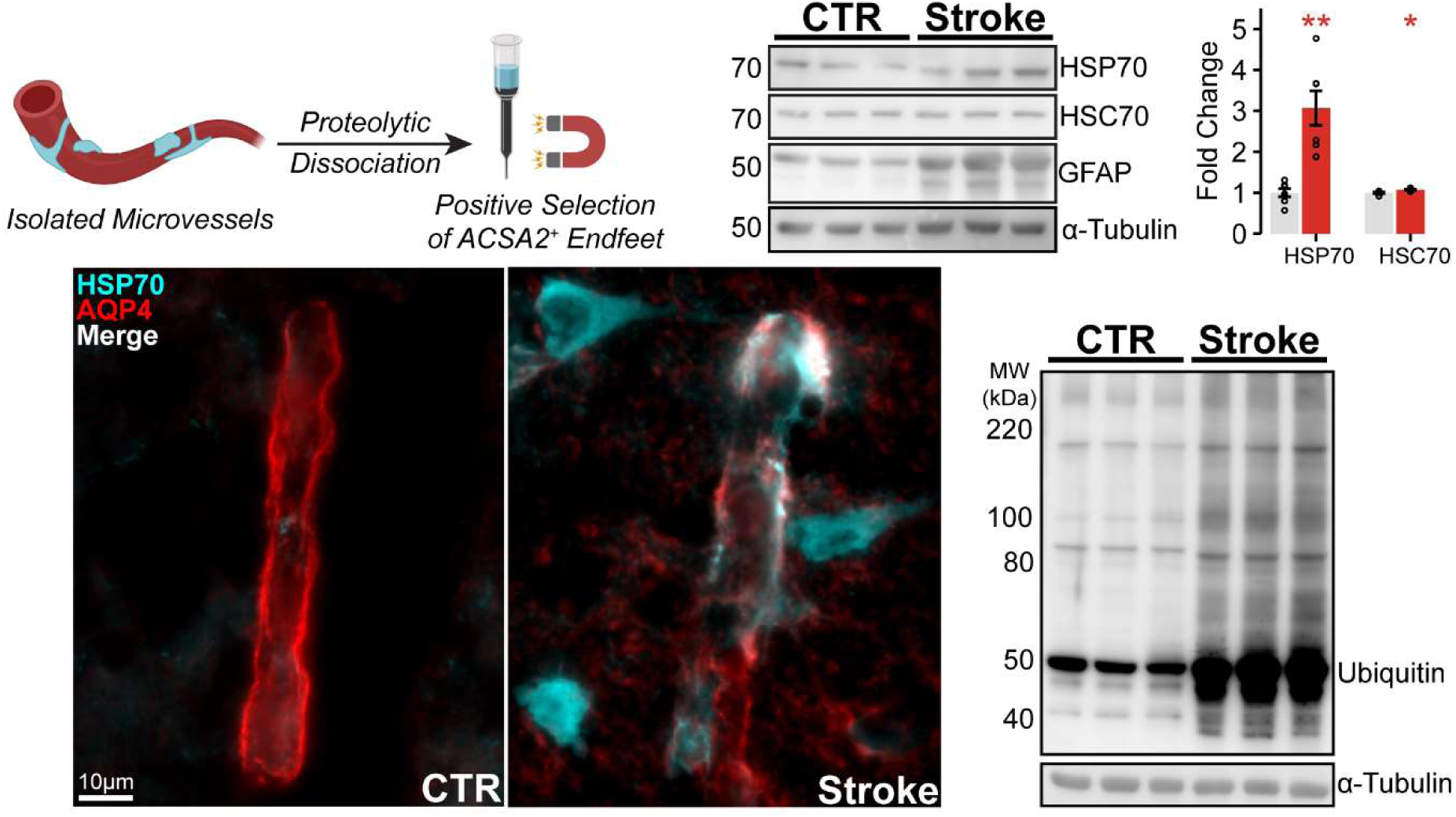
Upregulation of HSP70 and protein ubiquitination in post-ischemic astrocyte endfeet. (A) Schematic illustrating the enrichment of perivascular endfeet, which involves proteolytic dissociation of isolated microvessels followed by magnetic separation of ACSA2+ endfeet. (B) Representative immunoblots of HSP70, HSC70, GFAP, and the loading control α-tubulin from isolated ACSA2^+^ endfoot lysates. Densitometric quantification of HSP70 and HSC70 normalized to α-tubulin are plotted as mean ± SEM (*n* = 6 biological replicates). (C) Representative high-powered images of the post-ischemic cortex with immunolabelling for HSP70 (green) and AQP4 (red) (scale bar = 10µm). (D) Representative immunoblots of ubiquitin and the loading control α-tubulin from isolated ACSA2^+^ endfoot lysates. (\**p* < 0.05, \*\**p* < 0.01).

## Discussion

In this study, we report the translatomic response to ischemia in perivascular astrocyte endfeet. We have identified a set of 205 differentially expressed genes in the astrocyte endfoot translatome after a 2-hour ischemia / 6-hour reperfusion insult to territory of the middle cerebral artery. While dysfunction of local translation in astrocytes has been implicated in other pathologies such as leukoencephalopathy [44] and ALS [45], it has not been demonstrated previously in stroke.

The changes we have observed in the endfoot translatome overlap with, but are distinct from, those previously reported in the whole astrocyte translatome. One common feature is the robust activation of the heat shock response, which may be explained by disruptions in proteostasis, as indicated by increased protein ubiquitination in astrocyte endfeet after stroke. We have shown using various assays, including RiboTag RNA-sequencing, immunohistochemistry, qPCR, and immunoblotting that HSP70 is induced in astrocyte endfeet after cerebral ischemia/reperfusion.

After acute molecular stress – classically, heat, but also ischemia and hypoxia – the transcription factor HSF1 trimerizes and is translocated to the nucleus, where it activates a broad transcriptional program known collectively as the heat shock response [46–49]. The molecular chaperone HSP70 is a central part of this response and rapidly becomes one of the most highly expressed proteins in the cell after acute stress [50]. HSP70 is upregulated in neurons, astrocytes, and microglia after cerebral ischemia [51,52]. Consistent with our results, genes encoding HSP70 are upregulated in the whole astrocyte translatome at four hours [22], but not at 72 hours, after stroke [21]. Overexpression of HSP70 rescues cells exposed to ischemia from apoptosis [53] and decreases the size of the ischemic lesion in vivo [54]. Our findings suggest that the heat shock response in astrocyte endfeet is mediated, at least in part, by local translation. Cerebral ischemia induces a range of other changes in proteostasis, including the unfolded protein response in the endoplasmic reticulum and activation of autophagy [55]. An open question is how these pathways may be altered in the astrocyte endfeet, and what the implications are of such responses to injury and repair after ischemia.

We observed that the rise in HSP70 coincides with a substantial increase in ubiquitination. A widespread increase in ubiquitination in astrocyte endfeet after ischemia has not been reported previously, but our results are reminiscent of earlier findings of ubiquitination in post-synaptic densities after ischemia [56]. Our results suggest that ischemia induces local disruptions in proteostasis that lead to the activation of proteostasis pathways. These effects of ischemia are not unexpected, as many components of the proteostasis network, including the proteasome and a range of molecular chaperones, are ATP-dependent. Therefore, in the ATP-starved environment of ischemia, the maintenance of protein folding is impaired. As we have assayed the astrocyte endfeet 6 hours after reperfusion, we may be observing the response to damage accumulated during the 2-hour ischemic period. Whether this is salutary or a contributor to reperfusion injury remains to the be determined. Future studies on the time course of translatomic and proteomic changes may yield mechanistic insights by defining the sequence of proteostasis changes that occur after ischemia.

In addition to the prominent role of the heat shock response, our findings suggest other intriguing local response mechanisms. Pathway analysis suggests that these underpinnings center around four major functions: cell cycle/death, inflammation, metabolism, and proteostasis. The transcription factors whose targets are enriched among our differentially expressed genes are consistent with this pattern. We recently reported a significant calcium influx in astrocyte endfeet after ischemia. Consequently, we note that a significant number of calcium-regulated genes are differentially expressed in the endfoot translatome after stroke. The most upregulated calcium-responsive gene is *Thbs1*, which encodes the cell adhesion protein thrombospondin 1 (TSP1) that plays a complex role in angiogenesis [57,58]. *Itpkb*, the most downregulated calcium-response gene, encodes inositol trisphosphate-3 (IP3) kinase-B, which regulates the IP3 second messenger system with wide-ranging effects on cellular function. It has a putative role in neurodegenerative diseases [59,60], but its potential role in stroke has not been described. Other differentially expressed genes have also been implicated in angiogenesis. *Ptx3* encodes the acute phase protein pentraxin-3 important in the classical complement pathway that has been shown to promote neurogenesis and angiogenesis in rodent models of stroke [61,62] and to regulate the blood brain barrier [63]. *Sfrp5* encodes the secreted frizzled-related protein 5, a Wnt-signaling pathway antagonist and anti-inflammatory adipokine that has been shown to regulate angiogenesis [64,65].

Beyond highlighting specific genes, our findings also suggest potential mechanisms by which translatomic changes in the astrocyte endfeet might be regulated. The enrichment of specific transcription factor targets could reflect either transport of de novo synthesized mRNAs from the nucleus or the maintenance of selective pools of mRNA in the endfeet that the ribosome translates after a stimulus. Our analysis points towards possible mechanisms for such regulation, including basic features of the transcript and association with RNA binding proteins. Actively translating transcripts in perisynaptic astrocyte processes have been shown to have longer 5’-UTRs, coding regions, and 3’-UTRs than their somatic counterparts [17]. They also have decreased RNA secondary structure in their 3’-UTR, but similar GC content throughout. We find that some of these features are also associated with the post-ischemic endfoot translatome. In particular, genes upregulated and downregulated in response to ischemia/reperfusion differ in their length bias. They also differ from the unchanged translatome in terms of GC content. This suggests that some of the same features that help to regulate localization of transcripts in peripheral processes may play a role in guiding their response-specific translation. Similarly, the RNA-binding protein QKI which has been shown to regulate a large number of transcripts in whole astrocytes is specifically associated with a substantial minority of the genes downregulated in response to ischemia/reperfusion. QKI and other RNA-binding proteins may be important in governing mRNA transport to and ribosome binding in astrocyte endfeet.

One limitation of our study is the specificity of current isolation methods. It has been reported that perisynaptic astrocyte process preparations do not completely remove neurons [17]. Validation experiments of our approach suggest a relatively pure preparation of astrocyte endfeet (Figure 1). However, transcriptomics approaches are likely highly sensitive to contamination from cell bodies, which have a larger absolute abundance of mRNA. Our results, which show the expression of transcription factors and somatic markers (Table S4), suggest that some amount of somatic contamination may persist with these techniques. That we see a distinct translatome from whole astrocyte responses to ischemia argues that we are able to identify endfoot-specific signals. In addition, we find overall low expression of endothelial markers in our RiboTag data set. Finally, we have validated our observations about HSP70 using more targeted approaches. In particular, immunohistochemistry of coronal brain sections is independent of isolation techniques and provides strong validation for the localization of HSP70 in astrocyte endfeet after stroke.

Taken together, we demonstrate that early phases of cerebral ischemia and reperfusion induce a rapid change in the translatome of perivascular astrocyte endfeet. Given the critical role of astrocyte endfeet in maintaining the BBB and the significance of endfeet swelling after ischemic stroke, the regulation of the endfoot proteome is highly relevant to the pathophysiology of stroke. Future investigations of the endfoot translatome would help to resolve several remaining questions, including the time course of translatomic changes; the differences between the translatome and stored pools of mRNA in endfeet; and the mechanisms of mRNA transport to endfeet. We anticipate that such efforts to determine the regulation of the astrocyte endfoot translatome will yield new biomarkers and therapeutic targets for ischemic stroke.

## Author Contributions

Conceptualization, B.S., P.C., V.G., and J.M.S.; methodology, B.S., P.C.; validation, B.S., P.C., C.T., R.S., K.K.; formal analysis, B.S., P.C., V.G., J.M.S.; investigation, B.S., P.C., C.T.,R.S., N.T.; data curation, B.S., P.C., C.T., V.G.; writing – original draft preparation, B.S., P.C.; writing – review and editing, B.S., P.C., V.G., J.M.S.; visualization, B.S., P.C., V.G.; All authors have read and agreed to the published version of the manuscript.

## Institutional Review Board Statement

The study was approved by the Institutional Animal Care and Use Committee (protocol #0522007, approved August 23, 2022) of the University of Maryland School of Medicine (Baltimore, MD, USA).

## Data Availability Statement

All data presented in this study are available upon request from the corresponding author.

## Acknowledgments

We acknowledge Dr. Shilpa D. Kumar and the Confocal Microscopy Core, Center for Innovative Biomedical Resources (CIBR), University of Maryland School of Medicine (Baltimore, MD, USA), for their technical expertise with STED imaging. We thank Dr. Luke J. Tallon with the Maryland Genomics, Institute for Genome Sciences (IGS), University of Maryland School of Medicine (Baltimore, MD, USA) for the RNA sequencing.

---

**Table S1.** List of antibodies for immunoassays used in this study.

| **Antibodies** | **Manufacturer** | **Catalog #** |
| --- | --- | --- |
| ***Immunohistochemistry*** |  |  |
| Rabbit polyclonal anti-GFP | Cell Signaling Technology | 2555S |
| Guinea pig polyclonal anti-S100B | Synaptic Systems | 287004 |
| Rat monoclonal anti-CD31 (MEC13.3) | BD Biosciences | 550274 |
| Rabbit polyclonal anti-AQP4 | MilliporeSigma | ab3594 |
| Mouse monoclonal anti-AQP4 (4/18) | Santa Cruz | sc-32739 |
| Mouse monoclonal anti-HSP70 (3A3) | Santa Cruz | sc-32239 |
| Goat anti-guinea pig IgG Alexa Fluor 488 | ThermoFisher | A11073 |
| Donkey anti-rat IgG Alexa Fluor 555 | ThermoFisher | A78945 |
| Donkey anti-mouse IgG Alexa Fluor 488 | ThermoFisher | A21202 |
| Donkey anti-rabbit IgG Alexa Fluor 555 | ThermoFisher | A31572 |
| Donkey anti-rabbit STAR RED | Abberior | STRED-1002 |
| ***Immunoblotting*** |  |  |
| Rabbit polyclonal anti-GFP | Cell Signaling Technology | 2555S |
| Rabbit polyclonal anti-HSP70 | Cell Signaling Technology | 4872T |
| Rat monoclonal anti-HSC70 (1B5) | Santa Cruz | sc-59560 |
| Rabbit polyclonal anti-GFAP | Dako | z0334 |
| Mouse monoclonal anti-ubiquitin (P4D1) | Cell Signaling Technology | 3936 |
| Rabbit monoclonal anti-tubulin (11H10) | Cell Signaling Technology | 2125 |
| Goat anti-rat IgG, HRP conjugated | Cell Signaling Technology | 7077S |
| Horse anti-mouse IgG, HRP conjugated | Cell Signaling Technology | 7076S |
| Goat anti-rabbit IgG, HRP conjugated | Cell Signaling Technology | 7074S |

**Table S2.**
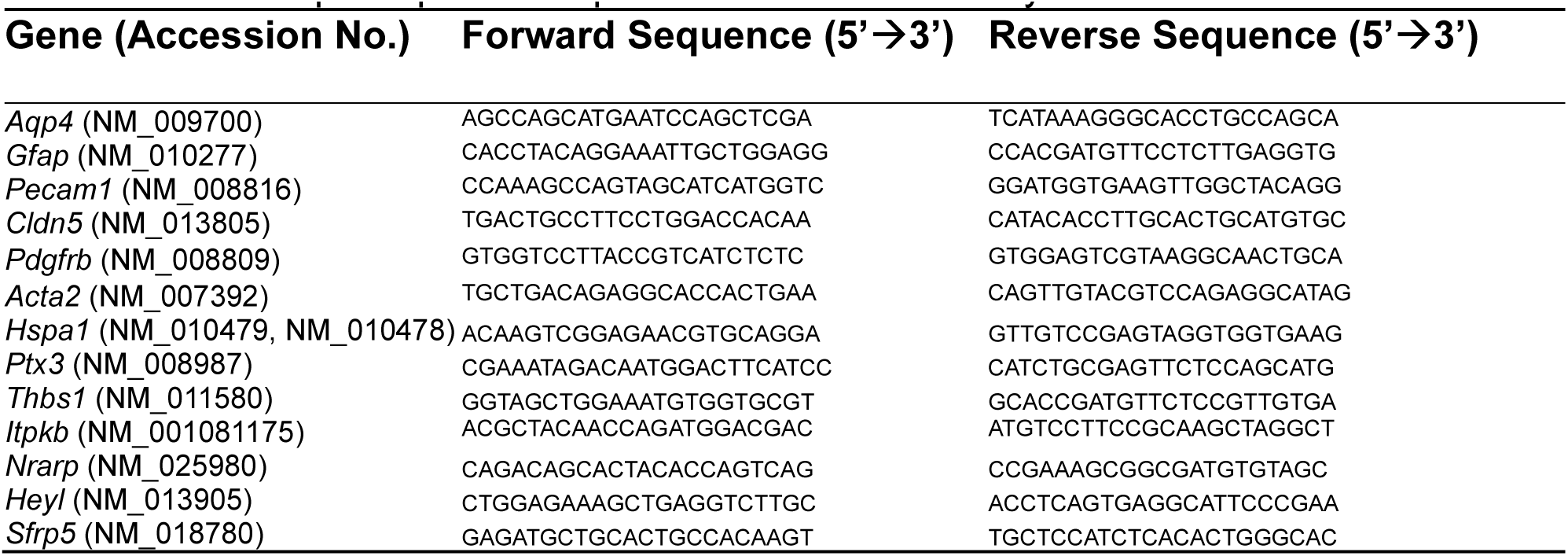
List of qPCR primer sequences used in this study.

